# Microphaser - a small-scale phasing approach for improved personalized neopeptidome creation

**DOI:** 10.1101/2021.08.11.455827

**Authors:** Jan Forster, David Lähnemann, Annette Paschen, Alexander Schramm, Martin Schuler, Johannes Köster

## Abstract

**Motivation:** Haplotype phasing approaches have been shown to improve accuracy of the search space of neoantigen prediction by determining if adjacent variants are located on the same chromosomal copy. However, the aneuploid nature of cancer cells as well as the admixture of different clones in tumor bulk sequencing data are challenging the current diploid based phasing algorithms. We present microphaser, a small-scale phasing approach to improve haplotyping variants in cancer samples. Microphaser aims to create a more accurate neopeptidome for downstream neoantigen prediction.

**Results:** Microphaser achieves large concordance with state-of-the-art phasing-aware neoantigen prediction pipeline neoepiscope, with differences in edge cases and an improved filtering step.

**Availability:** Microphaser is written in the Rust programming language. It is made available via Github (https://github.com/koesterlab/microphaser)and Bioconda. The corresponding prediction pipeline (https://github.com/snakemake-workflows/dna-seq-neoantigen-prediction) has been written within the Snakemake workflow management system and can be deployed as part of the snakemake-workflows project.

## Introduction

Cancer cells differ from healthy cells by accumulating various somatic variations, most importantly in protein-coding regions (González *et al.*, 2018). Those mutations arise from various sources such as single nucleotide variants (SNVs), short insertions and deletions (indels), gene fusions or larger structural variants like inversions, or duplications. Not all of those variants are necessarily cancer drivers, but still alter the genome, transcriptome and give rise to new mutated proteins, even if only acting as passenger mutations. Those neoproteins are cancer-specific and distinctively separate tumor cells from the surrounding healthy tissue (Schumacher *et al.*, 2019).

The degradation of intracellular proteins into smaller peptides - known as proteolysis or proteasomal cleavage - is an important process to remove flawed, misfolded proteins or to generate ligands for immune recognition (Calis *et al.*, 2015). To separate the shorter peptides from the complete proteome, we refer to the entirety of peptide sequences arising from proteasomal cleavage as peptidome. Cleaved peptides are then transported to the endoplasmatic reticulum and loaded on the major histocompatibility complex (MHC) proteins as peptide-MHC (pMHC) complexes (Neefjes *et al.*, 2011). Those complexes are presented on the cell surface and exposed to the T cells of the immune system. T cells have the capability to distinguish normal peptides from “foreign” neopeptides.

Therefore, the cancer-specific expression of those neopeptides can lead to an immune reaction similar to the reaction towards foreign pathogens such as bacteria or viruses, which can be exploited in immune response based cancer therapy (Schumacher and Schreiber, 2015),(Vormehr *et al.*, 2016). In immunology, an agent that is unknown to the host organism and will therefore elicit an immune response is known as an antigen. In analogy, immunogenic neopeptides which are derived from the patient’s cancer-specific genome are called neoantigens.

Tumor neoantigens are a key principle in tumor immunotherapy (the treatment of cancer by targeting tumor cells using the patient’s immune system). The inhibition of immune checkpoints such as PD-L1 or CTLA4 (immunomodulators which are overexpressed by tumor cells to downregulate immune reaction) is an already well established method used successfully in treating cancer patients (Pardoll, 2012).

Targeting of tumor neoantigens is a key principle in cancer immunotherapy. Different strategies are followed to mobilize neoantigen-specific CD8 T cells against tumor cells. Several approaches are based on the prediction of neoantigens from the tumor’s mutational landscape. Neopeptides derived from neoantigens are then used to design personalized vaccines.

The treatment outcome differs from patient to patient and needs to be assessed before individual therapy (Ribas and Wolchok, 2018). Since these approaches target the mutational landscape of tumor cells, there are useful markers for the effectiveness of immunotherapy such as the tumor mutational burden (Blank *et al.*, 2016), neoantigen burden (Schumacher *et al.*, 2019) or the diversity and expression of the HLA (human leukocyte antigen) antigen-presenting system (Boegel *et al.*, 2019). Precise in-silico prediction of neopeptides, their prevalence in the tumor and their antigenicity towards the immune system are crucial steps for optimising tumor immunotherapy. More personalized treatment options are currently being developed in the form of cancer vaccines (Sahin and Türeci, 2018), which rely on few accurately predicted and highly patient-specific neoantigen candidates.

It is not trivial to computationally predict valid neoantigen candidates from somatic variants for several reason. For once, not every SNV leads to an alteration of a protein. Not all emerging neopeptides can be bound and presented effectively by a patient’s MHC. Even if a neopeptide is a valid MHC ligand, it might still not elicit an immune reaction. Predicting the immunogenicity of an antigen is not yet completely solved, but can be estimated using a measure of (dis)similarity of a neopeptide towards the normal peptidome of a patient (Richman *et al.*, 2019). The difference in MHC affinity between the neopeptide and its normal equivalent can be a valid indicator for immunogenicity and is used in some existing workflows (Bjerregaard *et al.*, 2017). Low similarity and a strong increase in MHC affinity benefit the probability of a neoantigen to get recognised as “foreign” by immune cells, as the corresponding normal is either looking different or not well presented by the MHC at all. Those workflows focus on the comparison between the neopeptide and the normal peptide it was generated from.

However, neopeptides can also be similar to normal peptides arising from different genes and chromosomal regions. Thus, neopeptides should be filtered against the full normal peptidome of the patient, to reduce false positive neoantigen candidates.

Further, even a possibly immunogenic neoantigen might be too rare in the tumor to be a useful target for immunotherapy, either because it is only present in a small subclone or if it is not highly expressed.

Somatic variants such as SNVs should not be considered as isolated incidents but in relation to their haplotype. It has been shown that incorporating neighbouring variants - both germline and somatic - increases the success of neoantigen discovery (Hundal *et al.*, 2018; Wood *et al.*, 2020). If we link a neopeptide to a specific somatic variant that distinguishes it from its wildtype counterpart, it is evident that a second variant close to this somatic variant of interest can have an influence on the amino acid compositon of the resulting neopeptide. Even distal insertions or deletions can influence downstream variants by introducing reading frame shifts and completely changing the resulting protein. Therefore, in patient-specific neoantigen prediction, it is crucial to include all variants in order to create an as exact as possible search space (neopeptidome) to improve the success of downstream analysis. To model the possible co-occurrence of neighbouring variant sites, it is important to determine which variants co-occur on the same allele or chromosome copy. The combination of variants on a single chromosome copy is called a haplotype. The process to determine a haplotype from neighboring variants is called *phasing*. Phasing has already been incorporated in neoantigen discovery using methods like GATK’s ReadBackedPhasing (McKenna *et al.*, 2010) or HAPCut2 (Edge *et al.*, 2016). It has been shown that those methods successfully improve the accuracy of neoantigen prediction pipelines (Wood *et al.*, 2020).

However, all existing phasing approaches so far used for this purpose assume that the considered organism is diploid or at least of known ploidy and a homogeneous mixture of cells with the same genome. In cancer, these assumptions are usually not met. Here, we present microphaser, a tool for phasing possibly aneuploid sequencing data composed of different subclones, by looking at short and distinct sequence windows restricted to a length relevant for peptides.

## Approach

Microphaser is a small-scale phasing algorithm designed for resolving haplotypes of tumor data in short quasi-independent windows. Since correct genome-wide phasing is not yet solved for aneuploid tumor data, microphaser instead limits the problem to shorter, resolvable genomic regions. Since typical neoantigens are either 8-11 (MHC-I) or 15 (MHC-II) amino acids in length, determining the haplotypes of every possible neoantigen candidate independently is a reasonable simplification and a beneficial step in creating an exact neopeptidome. Microphaser uses a sliding window approach over the coding regions of the genome, while mapping genomic coordinates to gene annotations. Using a step-size of 3 (codon length), microphaser divides transcripts in window-sized peptides and determines the variant distribution at that position. Following the actual open reading frames (ORF), the resulting peptides represent all different haplotypes of the specific genomic region. On a paired tumor-normal sample, microphaser generates both a set of mutated neopeptides as well as their normal, unmutated counterparts. This allows the inclusion of a filter step to remove any mutated peptides which are already encoded by nonmutated sequences in the normal proteome.

## Materials and Methods

### Neoantigen prediction workflow: preprocessing

Microphaser can be run as part of a fully integrated neoantigen prediction pipeline which is realised using the Snakemake workflow management system (Köster and Rahmann, 2012). The pipeline is completely automated from start to end and self-contained in terms of software distribution and installation. The only exception are the netMHC-tools which are not available under a free license. All other needed software is installed automatically using the package management system conda, making use of the Bioconda channel (Grüning *et al.*, 2018). The pipeline operates on DNA-Seq data (whole-genome or whole-exome) in FASTQ format for matched tumor-normal samples. Optionally, RNA-Seq data can be provided for transcript abundance estimation. All required reference sequences and annotation are automatically downloaded from the Ensembl ftp-servers and cached by snakemake. Reads are aligned against the GRCh38 reference genome from the Ensembl release 100 using bwa-mem 0.7.17 (Li and Durbin, 2009), with subsequent sorting, indexing and deduplication using samtools 1.9 (Li *et al.*, 2009) and picard tools 2.22.1. Both somatic and germline SNVs and indels are called using the variant caller Strelka (Saunders *et al.*, 2012). Resulting germline and somatic variants are merged using bcftools 1.9 (Li, 2011), the germline-only calls however are also retained for use in the germline phasing step of microphaser. Variants are then preprocessed using bcftools norm, notably including left-aligning of indel variants and decomposition of multiallelic variant calls. Variants are annotated using the variant effect predictor VEP (McLaren *et al.*, 2016).

### Microphaser

For neopeptide identification, microphaser uses the tumor BAM file and the merged variant calls in BCF format, as well as a reference FASTA file and annotation in GTF format. Analogously, the generation of the sample-specific normal peptideome uses the normal sample BAM file and germline-only variant calls.

To reach maximal accuracy, microphaser tries to detect all possible haplotypes which carry somatic variation, including potentially small subclones which might exist in the tumor, leading to a representation of potential neoantigens in all cancer cell subpopulations.

The use of microphaser is structured in three main steps:

- creation of the normal sample-specific peptidome
- creation of the tumor-specific cancer peptidome
- translation of peptides and filtering for self-similarity

The creation of normal and tumor-specific peptidomes follows a nearly identical routine. In the following, we refer to the two cases as *somatic mode* and *germline mode*, and mention differences where necessary. First, genomic regions encoding for proteins are identified by defining transcripts and exonic regions per gene while parsing a genome annotation file. Microphaser then iterates over all valid transcripts exon by exon, using a sliding-window approach to identify all peptides of a specific length which might result from the current gene. Here, read-backed phasing is used to identify (sub)clonal haplotypes and their corresponding frequencies in the tumor sample. All resulting nucleotide sequences are then translated into amino acid sequence, while removing peptides resulting from synonymous mutation events by comparing them to their corresponding normal peptide. To further eliminate neopeptides which are similar to peptides arising from the normal peptidome of a patient—and will therefore most probably not be immunogenic—microphaser performs a filtering step against the complete normal peptidome while considering phased germline variants. The resulting valid neopeptides can then be evaluated in terms of differential MHC binding towards their respective corresponding normal peptides.

From the preprocessed data, microphaser starts reading gene, transcript and exon positions from the provided GTF (see Supplement) and produces phased peptides from both tumor and normal samples, following a sliding peptide-length window over the coding regions of a gene. The process of phasing is explained in the following section.

#### Phasing Algorithm

Each transcript of a gene is phased separately. Microphaser begins at the start position of the first exon and iterates over all exons of the transcript, before starting with the next transcript variant or gene. To this end, microphaser creates a peptide window with the length of the desired epitopes and slides the window over the transcript positions until the stop codon is reached. During phasing, microphaser follows the transcript’s open reading frame (ORF) encoded in the GTF to determine when a new codon has been added to the peptide sequence. However, indel variants might lead to a frameshift and create a new, additional ORF. Microphaser keeps track of all currently active reading frames.

The strand information of the specific transcript plays an important role in advancing the window throughout phasing. When reading genes located on the forward (3’) strand, the window advances with increasing genomic coordinates, sliding from “left to right”, with the window starting at the first position of the first CDS. Genes on the reverse (5’) strand are phased “backwards” (Fig. 2A). The start of the transcript is the end position of the first peptide window and the window slides from “right to left”. However, the sequence of the current peptide window is always generated from window start to window end, so from “left to right”. This is done to match the left-oriented notation of insertions and deletions in variant calls.

**Figure 1.**
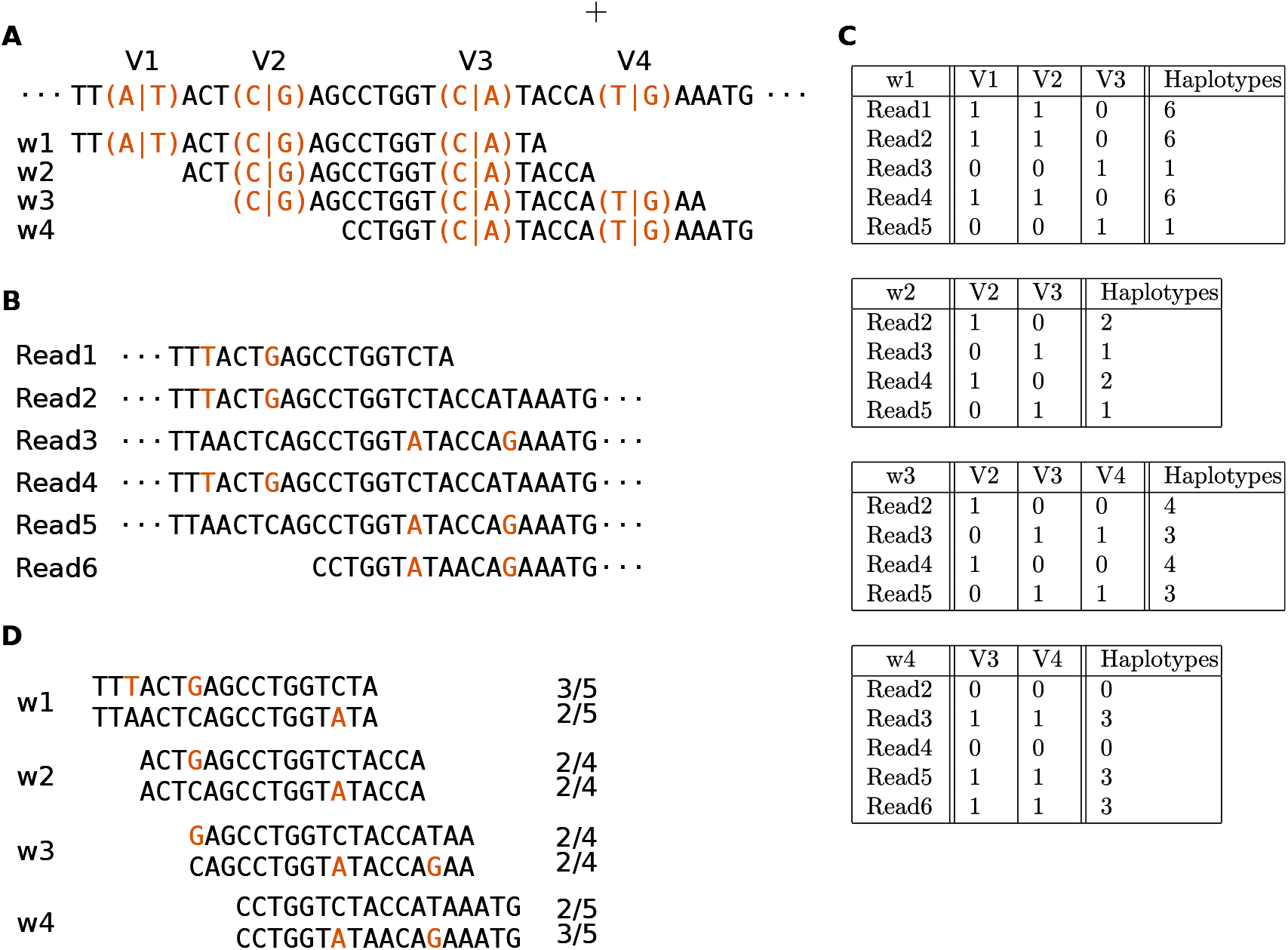
Example for small-scale phasing. A: Four SNVs embedded in the genomic sequence (top) and four resulting windows overlapping with the ORF. B: Six reads overlapping the genomic region in A. C: The ObservationMatrix changes representing all four windows in A. From window 1 to window 2, one read (Read1) and variant (V1) fall out of the scope. From window 2 to window 3, one variant (V4) is newly added. From window 3 to window 4, a new read (Read6) is added. D: The resulting haplotypes for all four windows, with frequencies computed per window.

**Figure 2.**
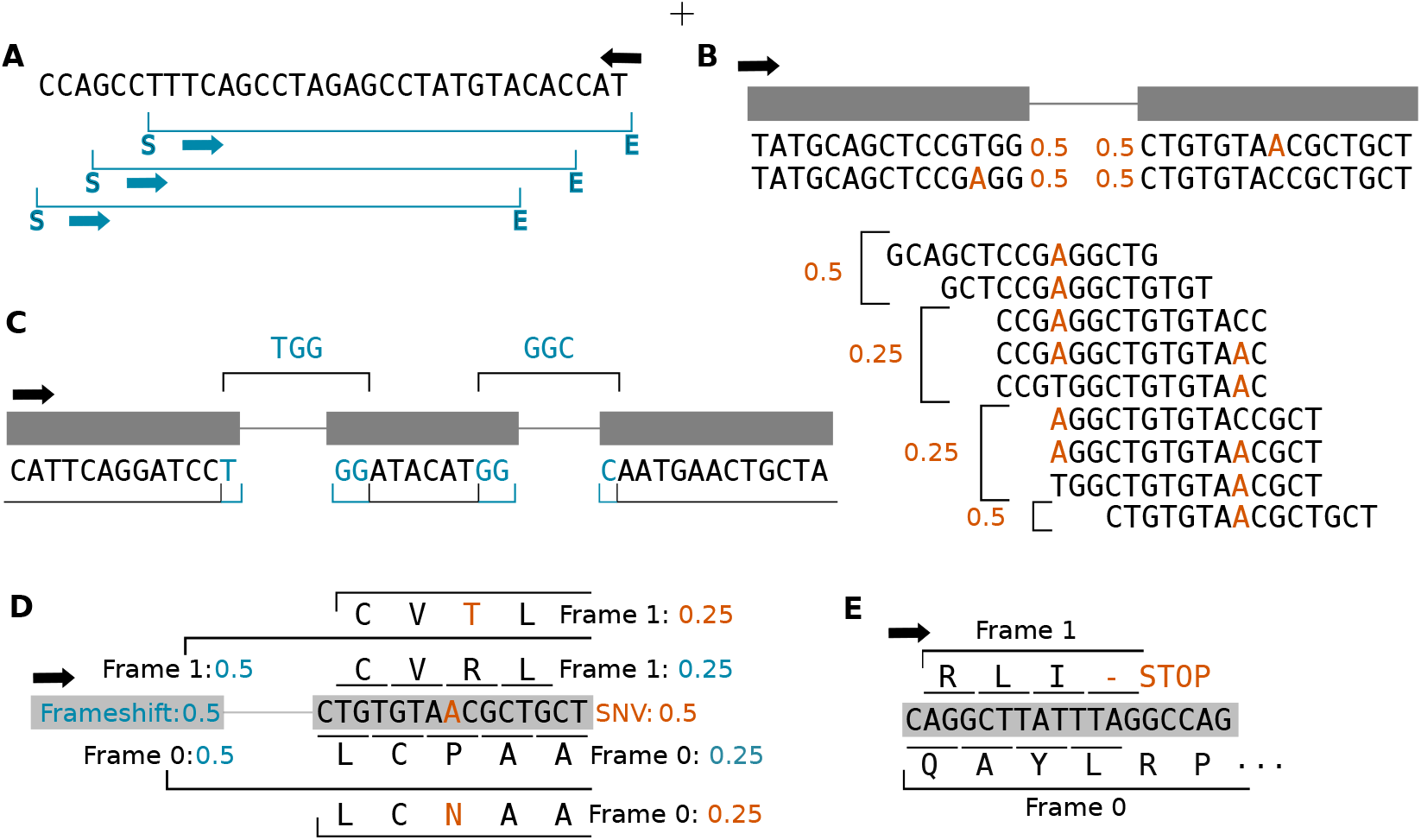
Algorithmic details. A: Phasing of reverse oriented transcripts with peptide windows. B: Joining of two exons at a splice-site with variants on both sides. The frequencies of the resulting neopeptides are computed from the combined exon window frequencies. Normal peptide fractions are not reported but can be implied from the neopeptide fractions C: Exon joining including an exon shorter than window size. The middle exon is completely integrated in the joining of the other two exon windows. Blue bases represent split codons. D: Representation of an SNV downstream of a frameshift variant. The frameshift is introduced in 50% of the reads and the DNA is transcribed in two ORFs with equal frequency. Downstream variants are integrated in both ORFs with respect to their frequencies. E: If a stop codon is reached in a window, the corresponding reading frame is finished and the analysis continues with the remaining ORF.

Microphaser uses read and variant information to determine every haplotype and its frequency in a peptide window. As the window advances, microphaser only considers reads and variants which are covered by the current window position, i.e. variants which lie inside the window and reads which completely enclose the window. To keep track of the variants and their supporting reads at every window position, the algorithm maintains an observation matrix. The observation matrix can be imagined as a bit-matrix M with dimensions *r*, the number of observations (or reads), and *v*, the number of variant sites, where every entry shows whether a read R supports the variant allele at site V as (Fig. 1C):

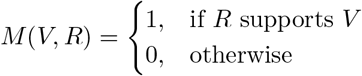

In this matrix, every read is associated to a bit vector with the size of the current variant list, where the n-th bit represents the n-th variant position inside the current peptide window. A bit is set to 1 if the corresponding variant is supported by the read, and set to 0 otherwise. This bit vector represents the subset of variants co-occuring in the read and therefore on the same haplotype. The representation of haplotypes as bit vectors allows the use of fast bitwise operations when adding and removing new variant sites.

The phasing itself can be written as a stepwise protocol:

#### Removing old reads

When advancing the peptide window, it is sufficient to consider the interval between the end of the previous and the end of the new window and remove reads whose end positions are located in that interval, in other words, observations which no longer span the current window.

#### Removing old variants

Similar to reads, variant sites can leave the current window after advancing. These variants are removed from the observation matrix by creating a bitmask of 2*^d^* − 1 with *d* representing the number of variants to be removed, using a bitwise AND operation between the bitmask and the haplotype of every read.

#### Adding new reads

Next, new reads are added to the matrix. For every read, an observation is initialized with a haplotype of 0. The haplotype is then updated for every variant site in the window by setting the corresponding bit to 1 if the read supports the variant.

#### Adding new variants

Finally, potential new variant sites are added. Again, microphaser only needs to consider the small interval between the end of the previous and the end of the current window. First, the haplotype of every observation is left-shifted by the number of added variants. Then, the haplotypes are updated considering their support of the newly added variants, analogous to step 3.

After every iteration of updating the observation matrix, microphaser checks if the window is in frame with the currently active ORFs (see start of section). Since the stepsize of the peptide window is one, this does at least happen every three iterations. In this case, all observations in the current observation matrix are grouped and every distinct haplotype is assigned with the number of supporting reads. For every haplotype, the reference sequence is updated by integrating the variants present in this haplotype—i.e., variants where the corresponding bit is set to 1. Somatic variants are only integrated in the neopeptide sequence (somatic mode), while normal variants are integrated in both normal peptides (normal mode) and neopeptides. After all variant sites in the window have been analysed, the mutated sequence is stored together with the normal peptide sequence. If a haplotype carries a somatic indel variant, the corresponding normal sequence is not reported since it is no longer similar to the mutated sequence (the normal sequence will however be stored during germline phasing). If a window does not contain any variant sites, or only germline variants, the sequence will only be considered in germline phasing mode. In the somatic mode, microphaser only writes peptide sequences containing somatic variants and their respective normal sequence. Neopeptides containing stop codons are not reported and since downstream regions after a stop will not be transcribed, the corresponding reading frame is removed from the list of active ORFs. For every valid neopeptide sequence, a record containing meta-information is created and saved under the same identifier (in a .tsv file). Important fields in this record are gene and transcript IDs, genomic position of the peptide window and the variant sites, haplotype frequency, read coverage and the DNA sequences of the mutated and the normal peptide. The haplotype information of the window is then stored for one additional iteration to ensure its accessibility in case of an exon joining event (see below).

#### Exon Gap Joining - Closing distances between variants

The segmentation of coding transcripts into exons is one limitation of working with genomic coordinates. Since the basic algorithm phases one exon after the other, it would fail to generate peptides stretching over exon junctions (splice-sites), and would miss potential neopeptides from somatic variants of interest within a window length from a splice-site (Fig 2C). Thus, when microphaser restarts phasing at a new exon, the last peptide window of the previous exon is combined with the first peptide window of the current exon to create a sequence of length 2*w* spanning the splice-site. Microphaser then iterates over the resulting sequence to generate all peptide-length sequences. Since there are no reads overlapping both parts of this splice-site sequence in whole exome or whole genome sequencing, microphaser does not perform the haplotype counting explained above, but reuses the haplotype information of the two merged windows, computing a mean coverage and combining variant sites as explained in the next section.

Additionally, an exon might end with an unfinished codon consisting of only one or two bases. In this case, the codon will be completed by the first one or two bases at the start of the next exon. Since microphaser follows the ORF, a naive implementation would ignore those split-codon bases on both sides of the splice-site since they are not in frame. To tackle this, microphaser increases the window length to cover those split-codon bases at exon end or start regions and evaluates possible variant sites at the respective positions.

Furthermore, some exons might be shorter than the window, meaning they will not encode for a peptide of sufficient length by themselves. For these short exons, the window length is extended towards subsequent exons until the required length is reached (Fig. 2C).

#### Combining distant variants

When handling splice-sites, one or even both windows might contain variant sites and therefore feature multiple haplotypes. Since we lack information by reads overlapping the intronic gap between the two exons, microphaser cannot count haplotypes directly but considers all combinations of haplotypes from the two windows upstream and downstream of the splice-site. Haplotype frequencies for the newly generated peptide sequences are computed from the frequencies of both input haplotypes treating them as independent variables. This approach follows the assumption of neutral tumor evolution of Williams *et al.* (2016). The model postulates cancer growth from a pre-cancer cell which already contains a substantial mutational burden, consisting of clonal (or public) variants. Mutations that occur later during tumor evolution are mostly subclonal (private) passenger mutations which do not experience selective pressure (neutral) and therefore grow with a steady allelic fraction that reflects their time of occurrence.

Therefore, even two variants with a similar subclonal allelic fraction did not necessarily co-evolve in the same subclone but were rather introduced at the same point in time. Accordingly, microphaser treats all variants as potentially independent events, as even if they occurred in the same subclonal cell, they could have occured on different haplotypes. The model tries to avoid making assumptions of co-occurrence of variants and rather seeks to broaden the search space without missing potential neopeptides. While the approach will lead to false positives, cases are rare and can be assessed when evaluating the final resulting neoantigen candidates - for example using transcriptome data, if available.

Let *w*_1_ and *w*_2_ be windows adjacent to the same intronic gap. In *w*_1_, a heterozygous variant *v* results in two haplotypes *h*_1_ and *h*′_1_, while *w*_2_ is homozygous with haplotype *h*_2_. The possible haplotypes over the intronic gap are therefore *h* = *h*_1_*h*_2_ and *h*′ = *h*′_1_*h*_2_. The frequencies of those two haplotypes are computed as follows:

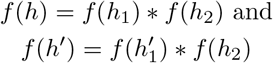

where *f*(*h*) is the frequency of haplotype *h*.

While iterating over the gap sequence, some windows will not include the variant site of *v*. In those windows, there is only one haplotype, and its frequency is computed as the sum of *f*(*h*) and *f*(*h*′).

Using this approach, microphaser also estimates the frequency of variants downstream of frameshift events. As mentioned above, all active reading frames are issued a specific frequency which is derived from the haplotype frequency of the frameshift variant. To estimate the probability of a downstream variant to be in phase with a reading frame, the ORF and variant frequencies are combined as described for exon gap joins.

### Keeping track of frameshifts

As mentioned in section, frameshift events (e.g., insertions or deletions) can lead to the generation of a second reading frame (heterozygous or subclonal frameshift) or even replace the original ORF (clonal or homozygous frameshift). Since these frameshift events do not only influence the peptide windows they are located in but the entire transcript downstream of the corresponding variant, it is important to keep track of the currently active reading frames. Therefore, microphaser defines every reading frame *F* as a tuple of the position of the corresponding frameshift variant, the introduced frameshift and its frequency *f*(*F*), derived from the variant’s allelic fraction. For all active ORFs *F* in a transcript, Σ*f*(*F*) = 1 holds.

For every transcript, microphaser starts with the main ORF *F*0, with a frequency *f*(*F*0) of 1. After the introduction of a frameshift by a variant, the frequency for the variant is computed and stored as the frequency of the new frame. Accordingly, the frequency of the original ORF is decreased by the alternative frame frequency.

From there on, the alternative frames are handled alongside the original frame. Downstream variants will be interpreted for all active frames, resulting in different neopeptides per frame. Especially variants which are synonymous with respect to the main ORF might generate neopeptides in the alternative frame.

If the alternative frame is flagged as somatic, all resulting peptides, even those not carrying SNVs themselves will be considered as neopeptides.

If an active reading frame reaches a stop codon, the frequency is set to 0 and the frame is removed from the set of active frames, since translation will terminate after the stop codon. However, any other active ORFs for which the codon does not translate to a stop will continue.

### Translation and Frequency Estimation

The phasing steps of microphaser produce peptide sequences still represented as forward oriented DNA bases. In the filter subcommand, these sequences are translated into amino acid (AA) sequences, depending on their respective transcript orientation.

After translation, mutated peptides which are identical to their normal counterparts and originate from synonymous variants are removed immediately. This is required since synonymous variants cannot simply be removed from the variant files as they can produce a non-synonymous AA sequence if they co-occur with other variants in the same codon or are located downstream of a frameshift. Furthermore, microphaser removes all peptides containing stop codons and does not translate peptides which are located downstream of a homozygous stop codon on the same ORF.

The remaining records are in most cases centered around somatic variants. Every somatic variant of interest results in k different peptides of length k. Assuming a standard epitope length of 9 AAs, one amino acid change generates 9 distinct neopeptides.

We strive to annotate the neopeptides with the frequency *θ* in which their underlying haplotype occurs at the given locus. This can be calculated in a Bayesian way as follows. Assuming that a haplotype is covered by *i* = 1, …, *k* peptide windows, our observed variables are *a_i_*, the number of reads supporting the variant haplotype and *n_i_*, the number of total reads over each window *i*. The true frequency *θ* of the variant haplotype is the latent variable we would like to infer. Then, the likelihood of *θ* can be calculated as

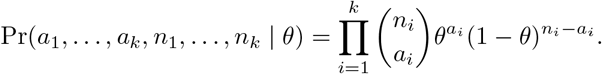

In other words, we assume that the observed counts are binomial distributed with success probability *θ*. Assuming a uniform prior distribution for *θ*, we can calculate the 95% credible interval and report it along with the maximum likelihood estimate for *θ*.

### Germline Phasing and Filtering

Not all non-synonymous variants necessarily lead to a true tumor-specific neopeptide. Considering the size of the human proteome, it is possible that a somatic mutation generates a peptide which differs from its direct corresponding normal peptide but still is identical to another normal peptide translated from a different gene. To detect those self-identical peptides, the germline mode in microphaser generates a patient-specific normal peptidome by incorporating and phasing only germline variants into the reference genome. These peptides are translated and stored in a hashmap serving as a control for neopeptide candidates. Any predicted neopeptides are compared against the entire normal peptidome to remove self-peptides.

### Neoantigen Prediction Workflow: Downstream Analysis

Output from microphaser can be used as part of a neoantigen prediction pipeline, using the resulting neopeptides as a candidate set or search space. In such a pipeline, the antigenicity of the resulting tumor neopeptides must be assessed with respect to the HLA alleles of the patient sample. In our workflow, HLA typing for class I HLA alleles is performed using optitype Szolek *et al.* (2014). The binding affinity of neopeptides, as well as their wildtype analogues, to the predicted HLA alleles is then obtained using netMHCpan v.4.1 Reynisson *et al.* (2020) for MHC-I alleles and netMHCIIpan v.4.0 Reynisson *et al.* (2020) for MHC-II alleles. The binding affinity is always predicted for a neopeptide and its corresponding normal peptide, in order to detect differential binding affinity due to the somatic AA change.

## Results

### Evaluation against neoepiscope

We compare the performance of microphaser with that of neoepiscope, a similar tool for neopeptide candidate generation from variants. Neoepiscope works on whole exome sequencing data and allows the use of pre-phased variant calls, with haplotype phasing performed by HAPCut2. Since neoepiscope does not provide a fully integrated pipeline, a snakemake workflow was created. Neoepiscope uses the allelic fraction information encoded in VCF4.1 format (https://samtools.github.io/hts-specs/VCFv4.1.pdf) to generate VAF information for the resulting neopeptides. HAPCut2 relies on genotype information for haplotype phasing. Therefore, STRELKA variant calls were updated to contain genotype information (GT) and allelic fraction (AF) fields using a pysam script. Further variant file preparations and haplotype phasing with HAPCut2 were performed as described in the neoepiscope best-practice. Neoantigen calling was used allowing non-canonical start and stop codons, with netMHCpan v.4.1 chosen for MHC binding affinity prediction. The complete neoepiscope snakemake workflow used in this paper is embedded in the complete analysis workflow and can be found archived on zenodo under https://doi.org/10.5281/zenodo.5163772, while a generalised version is available at https://github.com/jafors/snakemake-neoepiscope. Reference data in FASTA and GTF format was obtained from the Ensembl GRCh38 assembly, release 100.

### Datasets

We evaluated microphaser’s and neoepiscope’s perfomance on two datasets:

1. A concordance analyis was performed using a validation dataset consisting of 4 biological replicates from the same WGS sample. All replicates were produced from different sequencing protocols in different labs. Craig *et al.* (2016)
2. A dataset of four melanoma metastases from a single patient was used to demonstrate the algorithm on a typical use-case. This dataset has been analysed for neopeptides in a study by Schrörs *et al.* (2017).

#### Concordance Dataset

To assess the robustness of microphaser neoantigen predictions, we analysed concordance of its results across four replicates (EBI, Illumina, GSC, TGen) from the same tumor cell line (melanoma cell line COLO829), sequenced with different sequencing technologies in different institutes Craig *et al.* (2016) (https://ega-archive.org/datasets/EGAD00001002142). Since microphaser is supposed to work on the exon level, whole-genome reads were filtered to map to exonic regions. We ran the pipeline independently on each replicate and compared the resulting neoantigen candidates between replicates by their predicted frequency (Fig. 3D,E). Additionally, we computed mean frequencies over the predicted neoantigen frequencies of all four replicates and compared the results of each replicate with the computed mean values (Fig. 3A,B).

**Figure 3.**
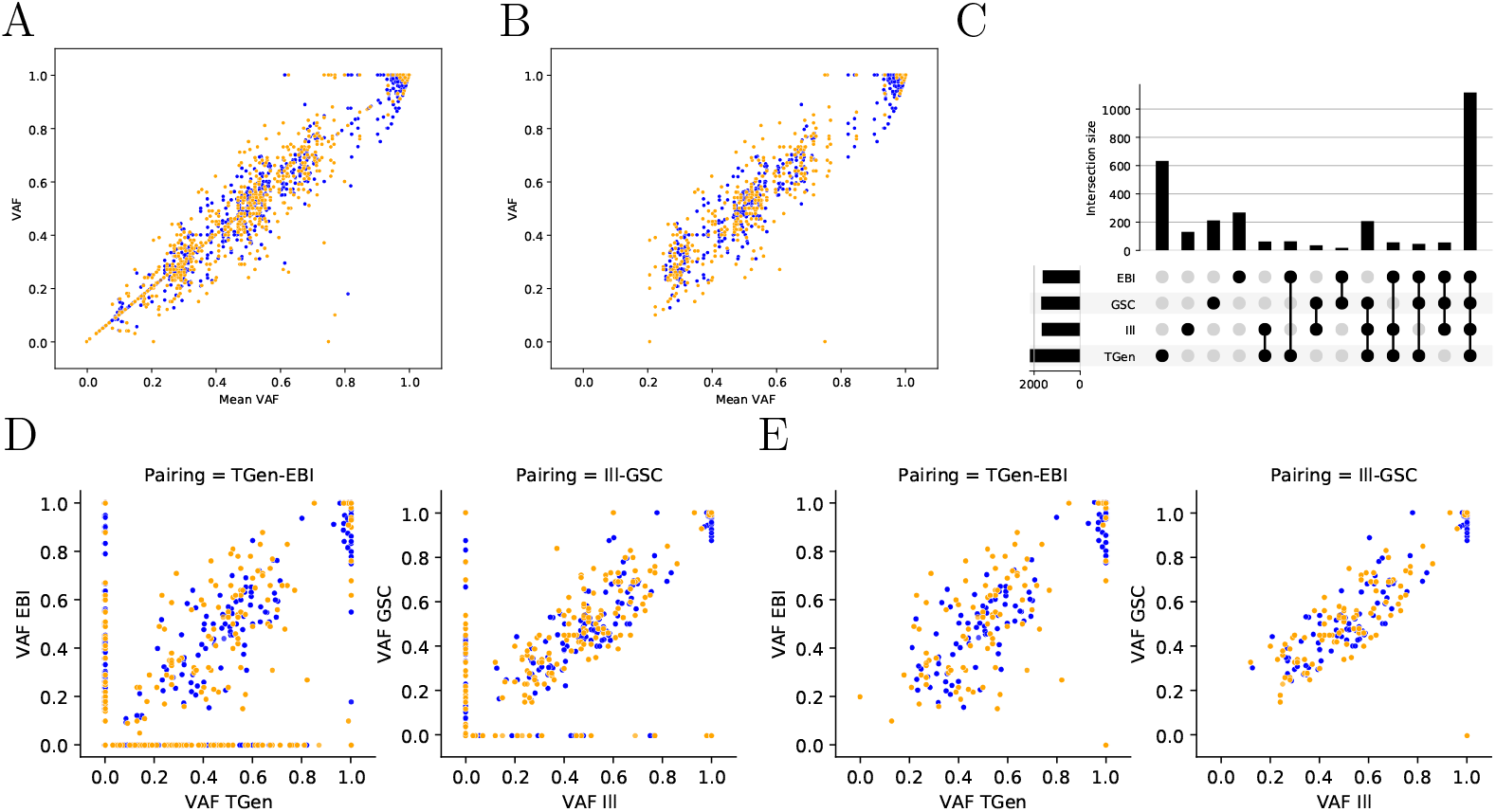
Concordance between four replicates of the same tumor-normal pair for both microphaser (orange) and neoepiscope (blue). A + B: Frequency for every neopeptide from a single replicate plotted against the mean frequency of the same neopeptide over all replicates (A: All variants, B: Variants shared by all replicates.) C: Upsetplot of all variants called in the four replicate pairs. D + E: Pairwise comparison of neopeptide frequency for two replicate pairs. A VAF of 0 represents a neopeptide not present in one of the compared replicates. (D: All variants; E: Variants shared by all replicates)

The same analysis was also performed using the neoepiscope pipeline (Section).

Microphaser shows good concordance between the replicates, similar to the concordance reached by neoepiscope (Fig. 3). Outliers and missing neoantigen candidates almost exclusively occur in regions with low coverage (¡ 15) and inappropriate sampling. Since most missed neopeptides probably result from differences in variant calling results across replicates, a second analysis was performed only on those variants shared by all four replicates. This removes most problematic neopeptides (zero vs non-zero VAF) from the comparison (see Fig. 3 D vs. E, and A vs. B). The remaining outliers can be explained with low coverage, low VAF and inadequate read sampling, where reads supporting the variant are not completely overlapping the peptide window.

In addition to the concordance analysis, we compared microphaser results directly with neoepiscope on all four replicates of the dataset. Here, it can be seen that differences arise mostly in neoantigen candidates predicted by neoepiscope which are not present in microphaser results (Fig. 4 A and B). The mechanisms and algorithmic differences leading to those missing neopeptides will be explained on examples in the following section.

**Figure 4.**
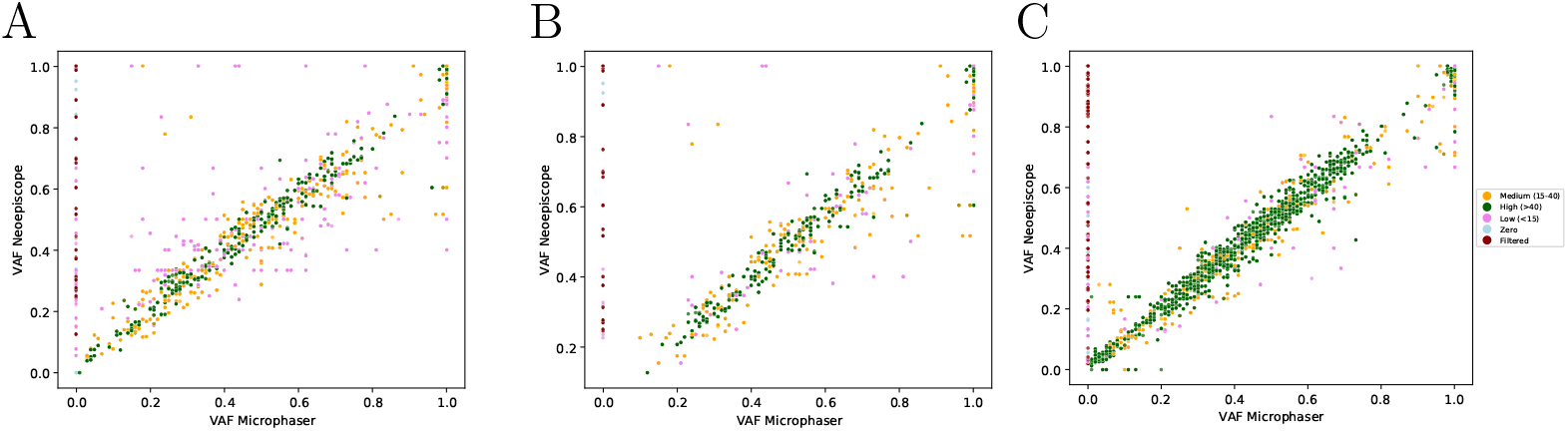
Comparison between neopeptides called by microphaser and neoepiscope. Frequency of neopeptides found only by one algorithm is set to zero in the other algorithm. (A: All variants; B: shared variants; C: Melanoma Metastases)

#### Melanoma Dataset

We used published WES data from four metastases of a single melanoma patient in order to test microphaser on a real case (Schrörs *et al.*, 2017). In this study, four neoantigen candidates were validated with T-cell assays, which could all be predicted by both microphaser and neoepiscope. This dataset was used to explore further differences in prediction results between microphaser and neoepiscope together with the aforementioned concordance dataset (Fig.4C).

### Comparison

In this section, we compare the neoantigen candidate sets predicted by microphaser and neoepiscope. We mainly focus on those neopeptides only predicted by either microphaser or neoepiscope and explain differences in prediction results due to algorithmic features and specific corner cases, illustrated on examples taken from the two analysed datasets.

#### Self-identity to Normal Peptidome

The lack of multiple neoepiscope candidates in microphaser predictions is the most prominent difference between microphaser and neoepiscope results and are highlighted in (Fig.4).

The largest fraction of such candidates not generated by microphaser is due to the filtering mechanism it uses. While neoepiscope only filters neopeptides where a synonymous peptide arises in the same transcript, microphaser filters neopeptides with synonymous peptides occurring anywhere in the complete normal peptidome of the patient. This especially leads to the elimination of false positive neopeptides due to homolog sequences in the same gene family. As can be seen in Fig. 5A, the S153L variant found in SIRPB1 which creates an altered amino acid sequence that is highly similar to a sequence in the gene product of SIRPG, a gene from the same gene family. All resulting 9-mers around the SIRPB1 variant are valid neopeptides compared to their position-equivalent normal peptide. However, comparing the 9-mer peptides of to the homolog sequence from SIRPG leads to the identification of four SIRPB1 tumor neopeptides identical to normal SIRPG peptides. These peptides are therefore removed from the candidate dataset during the microphaser filtering step.

**Figure 5.**
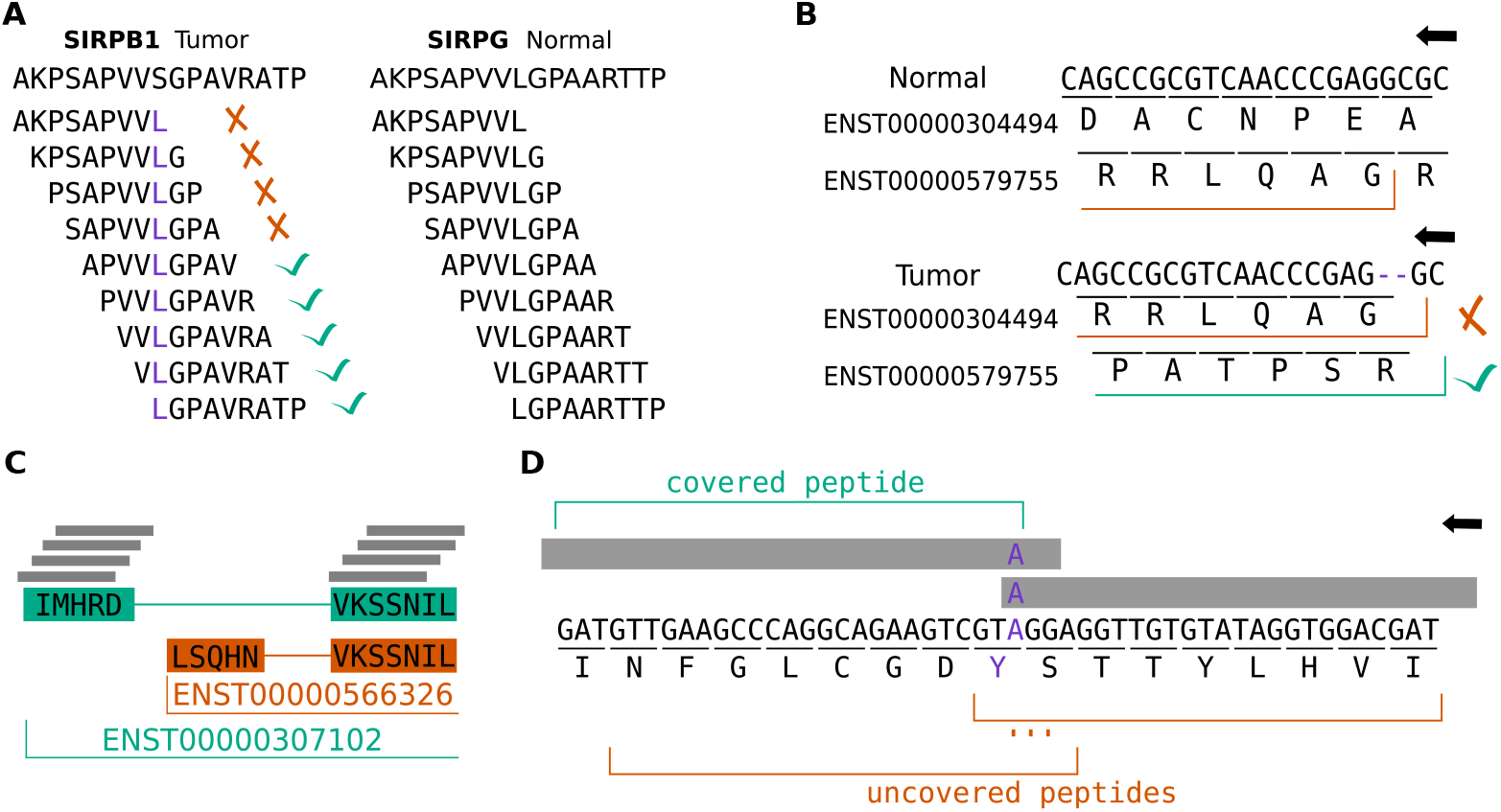
Interesting cases A: Filtering of identical peptides arising from non-synonymous variants due to similar genes. B: Two transcript variants of the same gene. A somatic deletion in transcript ENST00000304494 creates a new ORF which is identical to the normal transcript ENST00000579755 and will therefore be removed by microphaser. C: Varying read support of alternative transcript variants. One exon in the second transcript is not covered, which leads to uncertainty in the respective peptide windows. Microphaser will report a read coverage of 0. D: Influence of low coverage/VAF in PDZD7 gene. Only two reads carry the variant and their positioning does only allow to fully support one peptide window since read2 does not cover the entire mutated codon.

Furthermore, as seen in Fig. 5B, even a frameshift variant can lead to self-similar neopeptides. A deletion introduced in two transcript variants of CDKN2A generates a new frame for each transcript. However, the newly generated frame in the first transcript variant matches the original frame of the second transcript variant. Therefore, all peptides arising from this frameshift can be considered as self-similar. The second new frame has however not been seen in the normal transcript variants and does therefore count as a valid source of possible neoantigens.

#### Uncovered exons

Another source of differences in peptide prediction lies in regions with a lack of coverage in the WES. These cases especially originates from SNVs located near splice-sites. While the coverage at the variant site is fine—as the variant would not have been called otherwise—there is no read coverage at the next exon of the transcript (Fig. 5C). While neoepiscope uses the depth determined at the SNV site by the used variant caller, microphaser combines the read depth values of both exons at a splice site and returns a depth of zero if one of the exons has no coverage. In this case, the correct variant profile can not be determined—due to lack of read coverage—and it is likely that the resulting peptides might not be produced since the corresponding exon is not present in the patient. In Fig. 5C, two MAP2K1 transcripts are shown which share one exon with a P193S variant near the exon start. However, the previous exon is not the same in both transcripts. While the canonical transcript has a high read coverage on both exons, the alternative exon of the second transcript variant is not covered at all. Microphaser returns the resulting neopeptide sequences for both transcripts and indicates that the peptides unique to the alternative transcript have a coverage of zero, marking them as uncertain candidates.

#### Low Coverage

On the other hand, requiring that reads fully cover a window of interest to be considered for a contained variant, means that microphaser misses some peptides: when no read covers the entire peptide window, microphaser cannot resolve the haplotypes with absolute certainty (Fig. 5D). Therefore, microphaser is sensitive to regions with low coverage and small VAF. Especially if the reads which support the variant of interest are not entirely covering the genomic space around the variant, microphaser might miss some peptide windows. The impact of this sensitivity also depends on the length of the peptide window and the length of the reads. While the loss of single neopeptides is undesirable in terms of a complete overview of all neoantigen candidates, it can be argued that peptides resulting from sites with either low coverage, low VAF or both might not be ideal candidates since such sites carry an increased amount of uncertainty. However, this is a problem we would like to tackle in the future, e.g. by connecting reads which support the same haplotypes.

#### Long Deletions

Finally, longer deletions are not properly handled by the current microphaser approach in comparison to neoepiscope. Since the peptide windows are fixed at lengths specific to the desired peptide length, deletions in a window will shorten the peptide sequence in the window by the length of the deletion. Especially in large deletions, the windows will not contain enough bases to get a valid neoantigen out of it. However, long deletions could be treated as intronic gaps, with the windows before and after the deletions representing two neighbouring exons. We will explore this approach in the future.

#### Low Clonal Fraction

Another important source of differences lies in the phasing of distant variants, both germline and somatic. While the phasing of two distant SNVs - even located on different exons - is usually not crucial in detecting short neopeptides, there are two cases in which a correct phasing improves the prediction results:

1. If variants are located near the same splice-site on two consecutive exons, we want to know if they are on the same haplotype. However, since intronic regions are usually not covered in WES data, conventional read-backed phasing lacks reads to span the distance between the variants (see Fig. 2B). While microphaser computes a haplotype frequency for every possible combination of the two consecutive haplotypes, neoepiscope misses combinations and does not recompute the haplotype frequency but assigns the VAF of the somatic variant of interest predicted from variant calling to all neopeptides arising from this variant.
2. A special case appears with frameshift insertions and deletions. Often, it is not possible to determine which downstream variants are on the frameshift haplotype, especially considering the possibility of a shifted reading frame stretching over multiple exons. Here, microphaser also integrates all downstream variants in all current ORFs and recomputes the haplotype frequency based on ORF and variant frequency (Fig. 2D).

## Discussion

Microphaser is a tool for neopeptide candidate generation in neoantigen discovery, using small-scale read-backed phasing to determine haplotypes and haplotype frequencies, including variant combinations from small subclones. It performs phasing for both the germline and the somatic variants, generates all possible peptides based on the resulting haplotypes and extensively filters those to eliminate false positive neopeptides. By comprehensively eliminating neopeptide candidates with an identical peptide present in the normal peptidome of the patient (which are unlikely to elicit an immunogenic reaction), the filtering narrows the experimental search space for neoantigens.

We have shown that microphaser has a robust concordance over different conditions and replicates, especially in regions with a sufficient read coverage.

In comparison to other state-of-the-art phasing-aware toolkits such as neoepiscope, microphaser performs on a similar level and predicts a largely overlapping set of potential neoantigens (Fig. 3 and 4). However, microphaser explores neoantigen candidates more carefully, filters self-similar neopeptides comprehensively, and comes as part of a completely automated workflow incorporating all steps from read data to final neoantigen prediction. Furthermore, microphaser does not rely on published phasing tools such as HapCut2 or GATK ReadBackedPhasing, which are limited to diploid samples. Instead, it makes more accurate predictions, even for smaller subclones.

Due to its sliding window approach and the required read coverage of the complete window, microphaser finds fewer neoantigen candidates in low coverage regions than neoepiscope. While this enables microphaser to eliminate possible false positive peptides with partially zero coverage, it might also introduce false negatives at sites with a low variant allelic depth. Since the affected variants likely have a low read depth or allelic fraction, it can be argued that the resulting peptides are non-ideal candidates. But, higher read coverage per window could be achieved by using all reads which cover the variant site(s) in a window but not necessarily the complete window.

The filter step against the normal peptidome has been shown to successfully remove self-similar neopeptides.

While the normal peptidome is built patient-specific, future releases of microphaser will keep redundant peptides, i.e. peptides which do not arise by patient-specific germline variants but from regions identical to the reference peptidome, stored in a separate cache to avoid computational overhead.

We will further explore the possibility to not only remove synonymous neopeptides but also detect all normal peptides with a hamming distance of one. As seen in Section, MHC affinity is compared between a tumor neopeptide and its corresponding normal peptide (the same peptide without the somatic variant), since difference in affinity can serve as a predictive marker for immunogenicity. Usually, this is the normal peptide which is most similar to the neopeptide.

For neopeptides derived from insertions or deletions, this assumption does not hold, since the variant might introduce a large change in the normal peptide, including a potential frameshift. Finding all similar normal peptides to a neoantigen candidate might additionally improve the predictive value of the MHC affinity comparison between neopeptide and normal peptide. Therefore, we aim to improve the filtering algorithm to detect all normal peptides with a distinct Hamming distance towards every neoantigen candidate in order to increase the range of MHC affinity comparisons.

Considering subclonal samples, microphaser is able to resolve even small subclonal haplotypes. If a major clonal variant A is adjacent to a second subclonal variant site B, both a major clone A and a minor clone A-B exist. Microphaser detects both clones according to their frequency and returns peptides carrying only A and both A and B.

Another problematic case are longer deletions. In windows where deletions are introduced, microphaser has not yet analysed the genomic sequence downstream of the deletion and can therefore only remove the deleted bases from the sequence but not add further nucleotides after the deleted region. However, this case can be considered as a very short intronic gap and could be solved by a similar joining step, where the last complete peptide window before the deletion is joined with the first complete window after the deletion.

### 0.0.1 Future work

False negative results—especially in low coverage regions—arise from the strict requirement of reads overlapping the complete peptide window. We aim to combine information from overlapping reads supporting the somatic variant of interest into longer “pseudo-reads” which will negate the negative effect of unfortunate read sampling.

With the use of RNA sequencing data - which is important to analyse in neoantigen prediction to identify the expression level of potential neoantigens - we plan to identify a patient-specific transcriptome. Removing unexpressed transcripts and adding non-canonical transcript variants would very likely further improve the accuracy of microphaser. We aim to implement a phasing mode additionally or solely using RNA sequencing data in order to close intronic gaps and deliver a direct view on the expressed transcripts. Together with a variant calling approach detecting variation directly from RNA sequencing reads, this would allow for a purely RNA based neoantigen prediction. In this scenario, peptide frequencies would however no longer represent the haplotype frequency of the occurring variants but rather their fraction of the total expression level in this transcript.

